# HDAC2 inhibition restores H4K16ac and delays senescence in HGPS VSMCs

**DOI:** 10.1101/2025.11.07.687234

**Authors:** Rana Karimpour, Mzwanele Ngubo, William L. Stanford, Michael J. Hendzel

## Abstract

Histone H4 lysine 16 acetylation (H4K16ac) controls chromatin compaction, replication-coupled chromatin maturation and double-strand break repair, and is depleted in Hutchinson–Gilford progeria syndrome (HGPS) vascular smooth muscle cells (VSMCs). To identify the enzymes that set steady-state H4K16ac, we depleted KAT5, KAT8, all eleven zinc-dependent HDACs and all seven sirtuins in HeLa and U2OS cells, then in unaffected control and HGPS VSMCs. KAT8 depletion markedly reduced H4K16ac; KAT5 depletion had little effect. Among erasers, HDAC2 was dominant: its depletion or overexpression bidirectionally modulated H4K16ac, HDAC1 did neither, and SIRT1 depletion raised H4K16ac in these lines but far less in HGPS VSMCs. HDAC2-directed inhibitors (BRD4884, MI192, Santacruzamate A) restored H4K16ac in HGPS VSMCs, whereas the SIRT1 inhibitor EX527 did not. They also improved nuclear architecture, reduced γH2AX signaling, preserved Ki67 positivity across serial passage, and limited senescence-associated β-galactosidase accumulation. HDAC2 is therefore the predominant class I regulator of steady-state H4K16ac and a candidate target for limiting senescence in HGPS VSMCs.

## Introduction

Histone modifications shape the chromatin architecture that regulates transcription, DNA replication, and DNA damage sensing and repair ^1–4^. Histone H4 lysine 16 acetylation (H4K16ac) is highly conserved, and its cognate KAT5/8 acetyltransferase family has an unusually broad phylogenetic distribution ^5^. Unlike most histone modifications, H4K16ac directly modulates higher-order chromatin structure, limiting both the formation of compact 30-nm-like fibres and cross-fibre association in reconstituted nucleosomal arrays ^5,6^, an effect attributed to weakening of the interaction between the H4 N-terminal tail and the acidic patch of an adjacent nucleosome, although loss of H4K16ac does not detectably alter higher-order compaction in mouse embryonic stem cells ^7^. In mammals, H4K16 is acetylated predominantly by KAT8 (MOF/MYST1), a component of the male-specific lethal (MSL) and non-specific lethal (NSL) complexes ^8–10^. Although NSL acetylates H4K16 in vitro^8^, bulk H4K16ac in vivo is predominantly MSL-dependent ^9,10^.

H4K16ac has central functions in genome maintenance. It supports the normal spatiotemporal program of DNA replication, and in Drosophila MOF-dependent H4K16 hyperacetylation of the dosage-compensated X chromosome promotes replication-origin activation ^11,12^. Loss of MSL–KAT8 activity increases replication stress and DNA damage, and exacerbates chromosomal instability in cells with pre-existing genomic instability ^12–14^. H4K16ac also influences double-strand break repair pathway choice by antagonizing 53BP1 binding to H4K20me2-marked chromatin ^15,16^, thereby facilitating BRCA1 recruitment and homologous recombination ^16–19^.

In Saccharomyces cerevisiae, replicative aging reduces Sir2 protein abundance and increases H4K16ac ^20^. In mammals the direction is context-dependent, but reduced H4K16ac is the outcome reported in most systems examined to date. Loss has been observed in replicatively senescent WI-38 lung and BJ foreskin fibroblasts and in RAF- and RAS-induced senescence of WI-38 and IMR-90 fibroblasts ^21^, in replicatively senescent human diploid fibroblasts ^22^, and in human dermal fibroblasts aged in vitro ^23^. It has also been reported in aged mouse epidermis, oocytes, lung, haematopoietic stem cells, kidney and liver ^24–28^; aging human liver and progeroid Zmpste24−/− mice ^26^; and coronary artery vascular smooth muscle cells from older donors ^29^. The response is not uniform even among fibroblasts: in the same study, oncogene-induced senescence lowered H4K16ac in WI-38 and IMR-90 fibroblasts, which assemble prominent senescence-associated heterochromatin foci, but not in RAS-senescent BJ fibroblasts, which do not compact chromatin to the same extent ^21^.

Reported exceptions appear to depend on cell type, senescence programme, disease state, or genomic location. Of particular relevance, one study reported increased H4K16ac in HGPS dermal fibroblasts, which the authors attributed to impaired lamin A/C-dependent regulation of HDAC2 in progerin-expressing cells ^30,31^. Notably, lamin A/C binding was mapped to a *CDKN1A* promoter region previously shown to recruit HDAC2. In HGPS fibroblasts lamin A/C occupancy of that region increased while HDAC2 occupancy fell, despite higher total HDAC2 protein, suggesting impaired recruitment ^30^. Loss of lamin A/C– HDAC2 association correlated with elevated p21 ^30^. Oncogene-induced senescence increases H4K16ac in BJ fibroblasts, a response the authors report to be specific to that cell line rather than to the senescence programme ^21^; the same BJ cells lose H4K16ac during replicative senescence ^32^. H4K16ac also increases in fibrotic mouse and human lung but amidst a lower baseline H4K16ac in aged than in young mouse lung ^28^. Replicatively senescent fibroblasts show a two- to threefold increase in the number of H4K16ac peaks, concentrated at promoters of expressed genes, while being depleted of the mark across late-replicating regions and SAHF ^33^. Similarly, the number of H4K16ac peaks increases in the aging human brain but is lower in Alzheimer’s disease than in age-matched controls ^34^.

We previously found that loss of H4K16ac is pronounced in vascular smooth muscle cells (VSMCs) from Hutchinson–Gilford progeria syndrome (HGPS) ^29^. Classical HGPS is caused by a de novo LMNA mutation that activates a cryptic splice donor, producing progerin, an internally deleted prelamin A isoform lacking the ZMPSTE24 endoproteolytic cleavage site ^35–37^. Progerin retains a stably farnesylated carboxyl terminus that anchors it aberrantly at the nuclear membrane ^35,38^. The result is a segmental progeroid disorder that reproduces selected features of physiological aging ^39,40^. Transcriptomic profiling of patient fibroblasts identifies each of proteotoxic and oxidative stress, dysregulated DNA-repair programmes, and altered growth and inflammatory signalling recurring in 80–100% of patients ^41^. In HGPS VSMCs these abnormalities are accompanied by extracellular-matrix remodelling and atherosclerotic and vascular-aging signatures, and they intensify with passage ^29^. HGPS VSMCs derived from patient iPSCs accumulate progerin and show DNA damage and compromised viability under stress ^42^, and progeroid smooth muscle cells are selectively vulnerable to biomechanical strain ^43^. Death occurs at a mean age of approximately 13 years from cardiovascular causes ^39,40^. We recently found that H4K16ac is substantially reduced in HGPS VSMCs through both decreased acetylation and increased deacetylation, accompanied by DNA damage, premature senescence, and aberrant engagement of non-homologous end joining at newly replicated chromatin ^29^. Lonafarnib, the only approved therapy, lowers mortality risk and improves several disease-associated phenotypes, but the benefit is partial ^44–46^. This motivates our exploration of H4K16ac restoration as one candidate epigenetic target.

Steady-state acetylation is determined by the balance between acetyltransferase and deacetylase activities ^47^. KAT8 is the principal mammalian H4K16 acetyltransferase, and its depletion causes a global loss of H4K16ac ^9,10,48,49^, TIP60-complex activity has been implicated in, and in some experimental systems is required for, local H4K16ac at DNA breaks. ^16,19,50^. The enzymes responsible for H4K16 deacetylation are less clearly defined and have been attributed to both sirtuins and class I HDACs ^32,51–53^. SIRT1 preferentially deacetylates H4K16 and H3K9 among core histone residues in vitro, and RNAi-mediated depletion of SIRT1 hyperacetylates both residues in U2OS cells ^52^, although combined SIRT1/2 depletion produced little or no increase in H4K16ac in HeLa cells ^32^. SIRT2 has additionally been implicated in mitotic and senescence-associated regulation of H4K16ac^21,53^.

Among class I enzymes, combined disruption of HDAC1 and HDAC2 increases H4K16ac on newly replicated chromatin ^54^. Because the deposition-related acetylation of newly synthesized histone H4 is at K5 and K12 rather than K16 ^55^, H4K16ac detected on nascent chromatin is unlikely to be a deposition mark and instead reflects a replication-coupled or post-deposition modification. Proteomic capture of replication forks places the HDAC1/2-containing NuRD complex on nascent DNA in progerin-expressing cells ^29^, and HDAC2 is largely responsible for H4K16 deacetylation during mouse oocyte maturation ^56^.

Nevertheless, HDAC1 and HDAC2 show substantial functional redundancy in several somatic contexts ^57,58^, and their experimental separation has been further hindered by the limited isoform discrimination of most class I HDAC inhibitors ^59^. However, compounds whose additional targets differ can be used, so that HDAC2 is the only isoform the inhibitors engage at the concentrations used ^60,61^.

Here, we define the relative contributions of candidate acetyltransferases and deacetylases to steady-state H4K16ac in human cells, including unaffected control and HGPS VSMCs. We screened KAT5, KAT8, all eleven HDACs, and all seven sirtuins in HeLa and U2OS cells before testing the dominant activities in the VSMC models. KAT8 depletion robustly reduced H4K16ac in every cell line, whereas KAT5 depletion had little effect. Among class I HDACs, HDAC2 depletion produced the clearest increase in H4K16ac, while HDAC1 depletion had minimal effects. SIRT1 depletion increased H4K16ac to a similar extent as HDAC2 depletion in HeLa and U2OS cells but was markedly less effective in HGPS VSMCs. Critically, pharmacological inhibition directed against HDAC2, but not SIRT1, restored H4K16ac in HGPS VSMCs, attenuated senescence-associated phenotypes, and maintained the Ki67-positive cycling fraction during serial passage. These findings identify HDAC2 as a candidate epigenetic target for slowing cellular aging in HGPS.

## Results

### KAT8 maintains steady-state H4K16ac in HeLa and U2OS cells

Steady-state H4K16 acetylation has been attributed largely to KAT8, which catalyzes the bulk of this mark as part of the MSL complex ^9,48,49^, whereas KAT5/TIP60 has been linked to increased H4K16ac in contexts that promote homologous recombination ^50^, with separate TIP60-dependent mechanisms regulating 53BP1 through H2AK15 acetylation ^62^ and acetylating and activating ATM ^63^. H4K16 acetylation limits 53BP1 retention on damaged chromatin ^16^ and, when pre-existing in euchromatin, directs S-phase repair toward homologous recombination ^18^; consistent with this, its loss is associated with preferential engagement of non-homologous end-joining machinery on newly replicated chromatin in progerin-expressing VSMCs ^29^. To determine their relative contributions under routine growth, we perturbed each enzyme and quantified H4K16ac. In HeLa cells, siRNA-mediated KAT8 depletion reduced per-nucleus H4K16ac integrated intensity by 60% relative to the non-targeting control, whereas KAT5 depletion with siRNA produced no detectable change (Figure 1A). Immunoblotting of acid-extracted histones confirmed this effect, with KAT8 depletion reducing H4K16ac by more than 70% relative to the non-targeting control, whereas KAT5 depletion had little effect (Figure 1B). Lamin A/C siRNA, included as an assay-positive control, reduced H4K16ac by approximately 50%. Immunoblotting of whole-cell lysates confirmed efficient target depletion, with >80% reduction in protein abundance relative to the non-targeting control, as determined by densitometry after normalization to the loading control (Figure S1A). Consistent with these results, KAT8 depletion reduced H4K16ac in U2OS cells, whereas KAT5 depletion produced only a slight reduction (Figure S1B). Together, these data support KAT8 as the predominant acetyltransferase maintaining steady-state H4K16ac under basal growth conditions in these cell lines.

**Figure 1.**
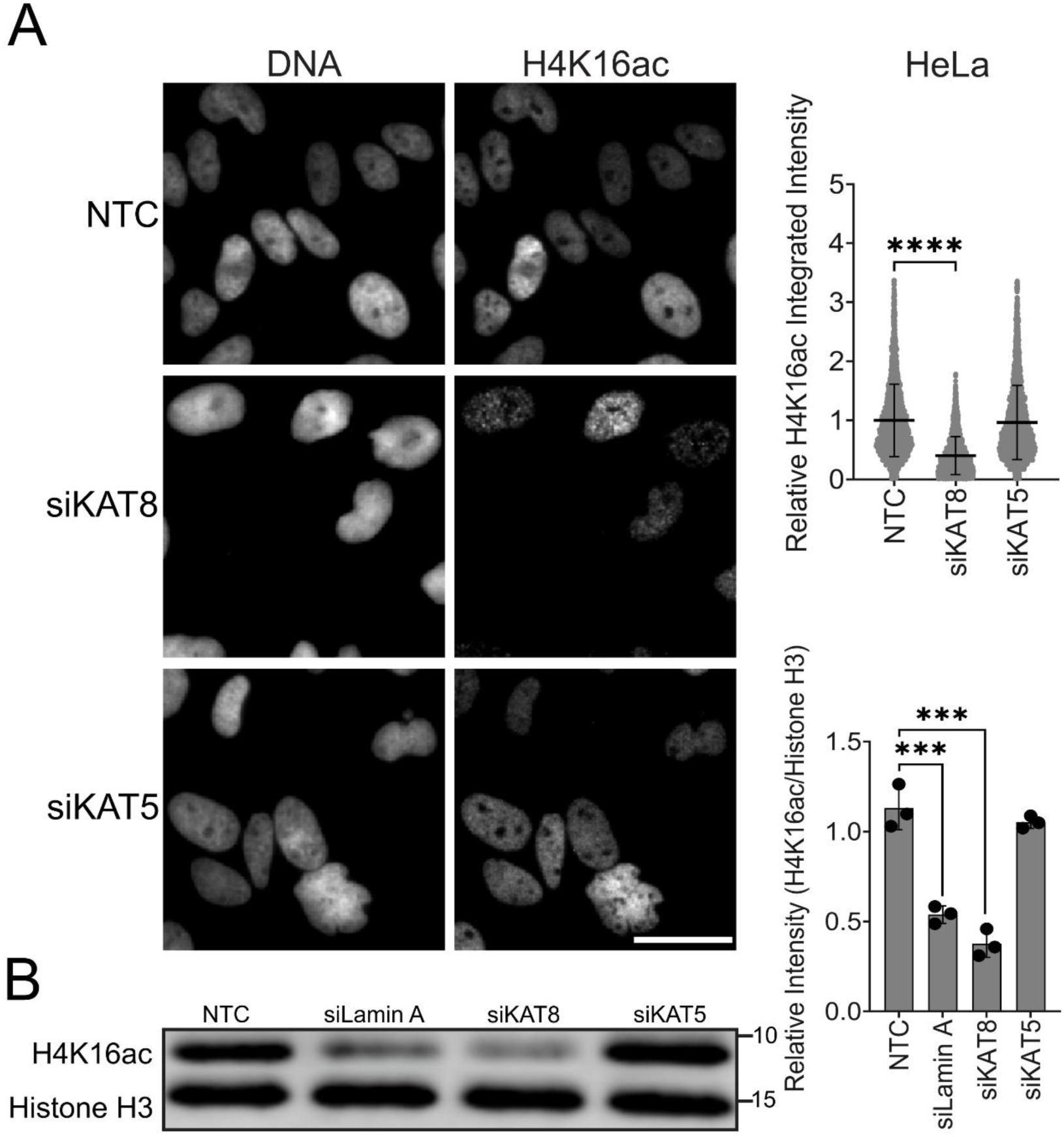
KAT8 maintains steady-state H4K16ac in HeLa and U2OS cells. (A) Representative immunofluorescence images of HeLa cells transfected with siNTC, siKAT8, or siKAT5, and single-cell quantification of nuclear H4K16ac integrated intensity. Points show individual cells; bars show the mean ± SD of three biological-replicate means, with approximately 1,000 cells analyzed per condition per replicate. RM one-way ANOVA, Dunnett’s vs. NTC. Scale bar, 20 µm. (B) Representative immunoblot analysis of acid-extracted histones from HeLa cells transfected with NTC, siLamin A, siKAT8, or siKAT5. Each point represents one independent biological replicate, and bars represent the mean ± SD from three separate biological replicates. RM one-way ANOVA, Dunnett’s vs. NTC. ***p < 0.001; ****p < 0.0001.

### HDAC2 is the predominant class I regulator of steady-state H4K16

Steady-state H4K16 acetylation results from a dynamic equilibrium between acetylation and deacetylation. To identify the relative contributions of individual deacetylases, we depleted HDAC1–HDAC11 and SIRT1–SIRT7 in HeLa and U2OS cell lines and quantified nuclear H4K16ac by immunofluorescence. HDAC2 knockdown increased mean per-nucleus H4K16ac by 42% in HeLa and 47% in U2OS cell lines relative to the non-targeting control (Figure 2A, S2A). SIRT1 knockdown also increased H4K16ac, by 47% in HeLa and 49% in U2OS cells. In contrast, HDAC1 knockdown had little effect in either line, despite the frequent association of HDAC1 and HDAC2 within shared corepressor complexes. These results distinguish HDAC2 from HDAC1 as the principal class I activity affecting steady-state H4K16ac under these conditions.

**Figure 2.**
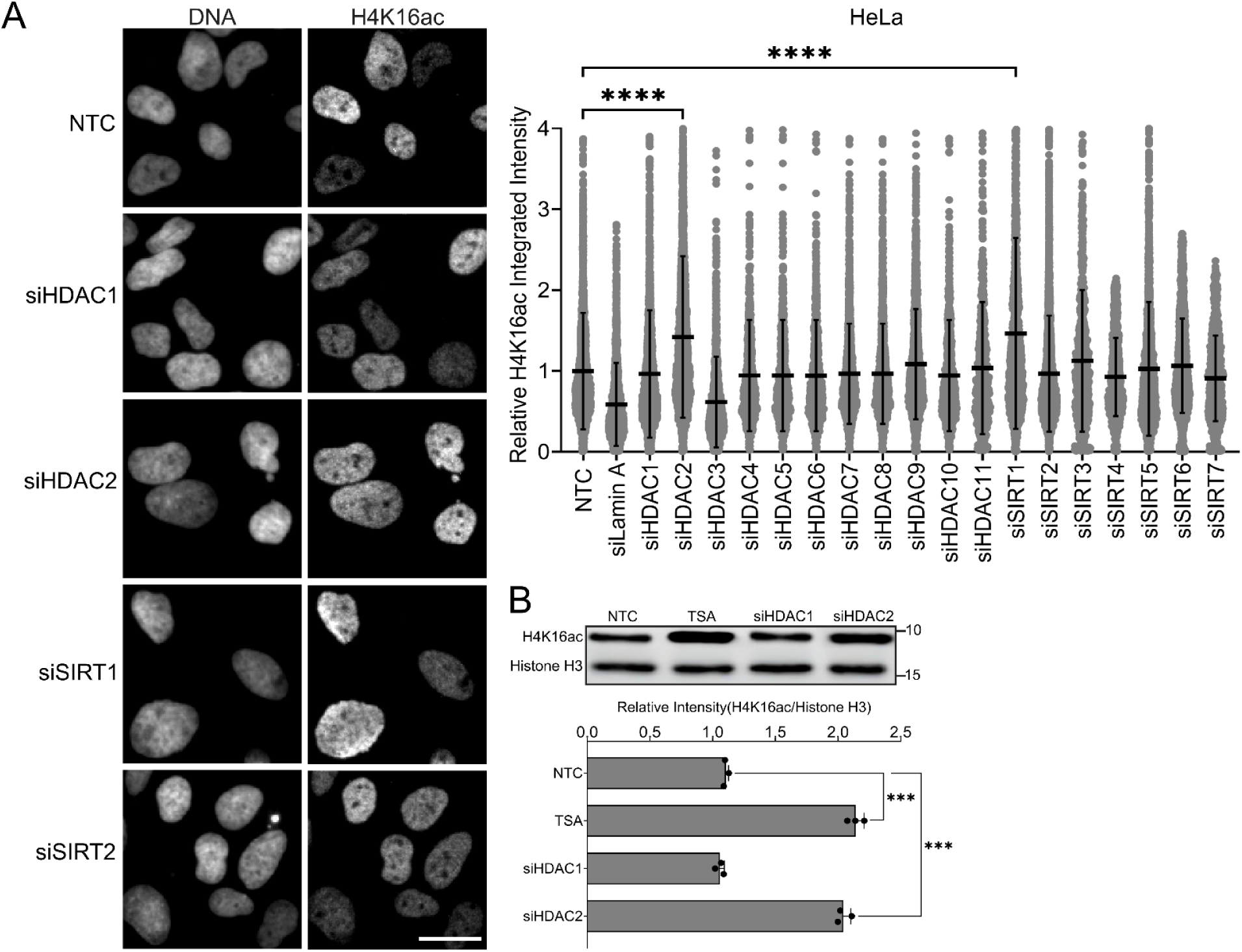
HDAC2 is the principal class I H4K16 deacetylase and a prominent source of cellular H4K16 deacetylase activity. (A) HeLa cells transfected with siNTC or siRNAs targeting Lamin A/C, HDAC1–11, or SIRT1–7, with representative images shown for NTC, siHDAC1, siHDAC2, siSIRT1, and siSIRT2 and single-cell quantification of nuclear H4K16ac integrated intensity. Grey points show individual cells; bars show the mean ± SD of three biological-replicate means, with at least 500 cells analyzed per condition per replicate. RM one-way ANOVA, Dunnett’s vs. NTC. Scale bar, 20 µm. (B) Representative immunoblot analysis of acid-extracted histones from HeLa cells transfected with NTC, siHDAC1, or siHDAC2, or treated with trichostatin A (TSA; positive control). Each point represents one independent biological replicate, and bars represent the mean ± SD from three independent biological replicates. RM one-way ANOVA, Dunnett’s vs. NTC. ***p < 0.001; ****p < 0.0001.

HDAC3 depletion produced discordant results between the imaging and biochemical assays. H4K16ac fluorescence decreased following HDAC3 depletion, whereas immunoblotting of acid-extracted histones from HeLa cells showed no significant reduction in bulk H4K16ac (Figure S2B). TSA, a potent inhibitor of class I and II HDACs ^64^, increased H4K16ac as expected (Figures 2B and S2B). These data indicate that HDAC3 depletion does not reproducibly reduce bulk histone H4K16 acetylation under these conditions and further distinguish HDAC3 from the robust H4K16 deacetylase activity associated with HDAC2. Knockdown of each of the remaining HDACs produced small changes that were not statistically significant.

Immunoblotting of whole-cell lysates confirmed efficient depletion of SIRT1, HDAC2, HDAC1, HDAC3, and lamin A/C, with residual protein levels of approximately 10–25% of the corresponding non-targeting controls (Figure S2C). Together, these loss-of-function experiments identify HDAC2 as the predominant class I regulator of steady-state H4K16ac in HeLa and U2OS cells, while also revealing a substantial contribution from SIRT1.

### Overexpression confirms HDAC2 is a robust H4K16 deacetylase

To complement the loss-of-function screen and test whether increased deacetylase abundance was sufficient to alter H4K16ac, cells were transfected with EGFP-tagged HDAC1, HDAC2, HDAC3, or SIRT1. In HeLa cells, HDAC2 overexpression reduced mean H4K16ac to 64% of the EGFP-only control, and SIRT1 overexpression reduced it to 66%, whereas HDAC1 and HDAC3 remained near control levels at 97% and 99%, respectively (Figure 3). U2OS cells showed the same pattern, with reduced H4K16ac following HDAC2 or SIRT1 overexpression but little change following HDAC1 or HDAC3 overexpression (Figure S3). Together with the depletion experiments, these reciprocal gain- and loss-of-function data support HDAC2 as the predominant class I regulator of steady-state H4K16ac in these cell lines and identify SIRT1 as an additional H4K16ac-regulating activity.

**Figure 3.**
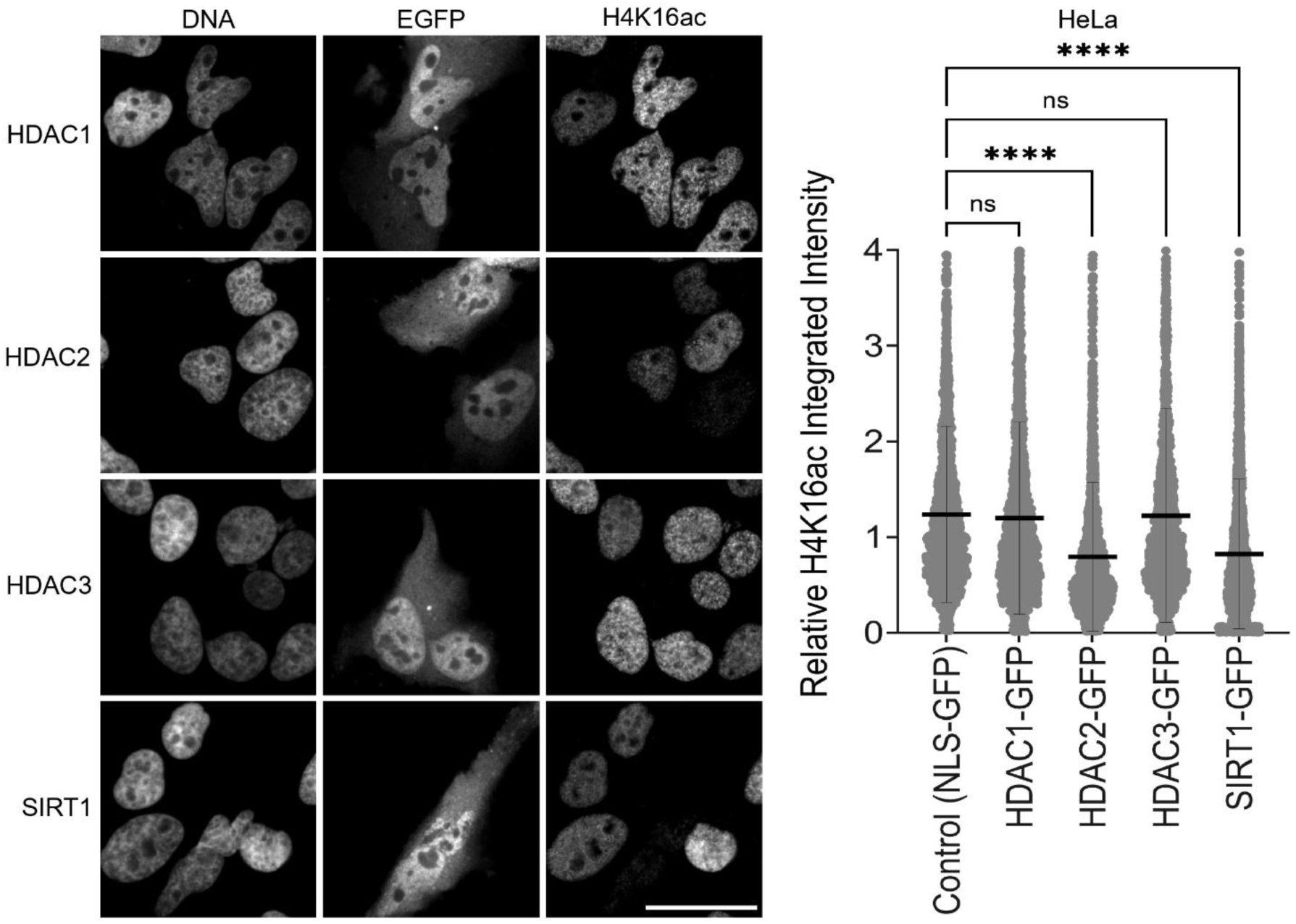
Overexpression confirms HDAC2 is a robust H4K16 deacetylase. HeLa cells transfected with EGFP alone or EGFP-tagged HDAC1, HDAC2, HDAC3, or SIRT1, with representative images and quantification of nuclear H4K16ac integrated intensity in EGFP-positive cells. Grey points show individual cells; bars show the mean ± SD of three biological-replicate means. RM one-way ANOVA, Dunnett’s vs. the EGFP-only control. ns, not significant; ****p < 0.0001. Scale bar, 10 µm.

### An HDAC inhibitor panel confirms HDAC2 as a druggable target for restoration of H4K16 acetylation levels

If HDAC2, but not HDAC1, is responsible for the increase of K16 acetylation upon treatment with class I HDAC inhibitors, preferential inhibition of HDAC2 might avoid some of the toxicities associated with general class I HDAC inhibitors ^65–68^, although these studies used pan-HDAC agents and do not attribute the observed toxicities to individual isoforms. We tested a panel of HDAC and sirtuin inhibitors to determine their influence on histone H4K16 acetylation (see Pharmacological Treatments). In particular, we made use of drugs that differ in their inhibition of individual members of the class I HDAC family in an effort to establish whether HDAC2 inhibition is sufficient to increase H4K16 acetylation abundance. We used BRD4884, an HDAC1/2 inhibitor ^61,69^; MI192, a low-nanomolar inhibitor of both HDAC2 and HDAC3 ^60^; Santacruzamate A (SCA), originally reported to be a highly potent HDAC2 inhibitor with only minor HDAC4 and HDAC6 ^70^ inhibition; Quisinostat (QS), a broad-spectrum HDAC inhibitor with potent class I activity ^71^; and EX527, a selective SIRT1 inhibitor ^72^. BRD4884 and MI192 have different reported class I profiles, HDAC1/2 and HDAC2/3 respectively, so HDAC2 is the principal class I activity common to both.

We titrated each compound in HeLa cells to choose working doses that produced clear increases in H4K16ac without growth inhibition or cell death (Figure S4A). The response to SCA was non-monotonic: H4K16ac increased maximally at 10 nM and diminished at higher concentrations (Figure S4A). At optimal concentrations in HeLa cells, SCA increased the mean H4K16ac by 62% relative to DMSO, MI192 by 34%, BRD4884 by 22%, and QS by 20%; however, the SIRT1 inhibitor EX527 showed no significant increase in H4K16ac (Figure 4). The strongest effects came from the HDAC2-focused inhibitors, consistent with HDAC2 acting as the primary H4K16 deacetylase under these conditions. Broader class I inhibition with QS gave a smaller increase. We repeated these experiments in U2OS cells. U2OS cells treated with the same inhibitors showed comparable responses, with increases of 62% for SCA, 34% for MI192, 23% for BRD4884, and 21% for QS, while EX527 showed no significant increase in H4K16ac (Figure S4B). The agreement across cell lines supports a model in which HDAC2 is the dominant contributor to H4K16 deacetylation, whereas EX527 did not phenocopy SIRT1 depletion under these conditions.

**Figure 4.**
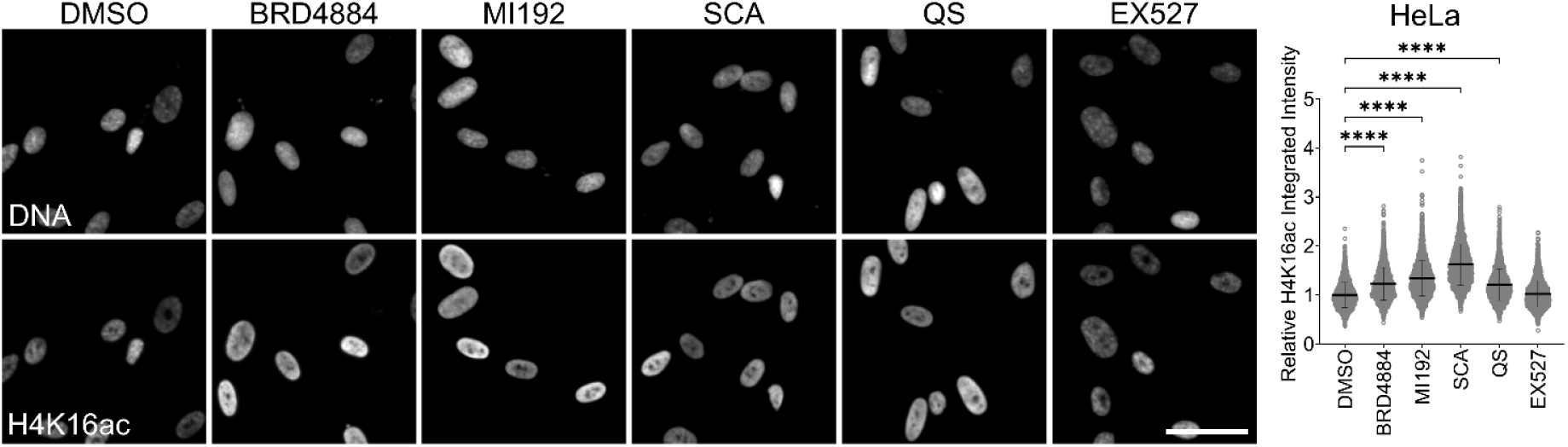
An HDAC inhibitor panel confirms HDAC2 as a druggable target for restoration of H4K16 acetylation levels. HeLa cells treated for 24 h with vehicle control (DMSO), BRD4884 (2 µM), MI192 (1 µM), Santacruzamate A (SCA; 10 nM), Quisinostat (QS; 250 nM), or EX527 (10 µM), with representative images and single-cell quantification of nuclear H4K16ac integrated intensity. Bars show the mean ± SD of three biological-replicate means. RM one-way ANOVA, Dunnett’s vs. DMSO. ****p < 0.0001. Scale bar, 20 µm.

### HDAC2 inhibition restores H4K16ac in HGPS VSMCs

HGPS VSMCs display reduced H4K16ac that is associated with more rapid deacetylation rates, increased NuRD association with newly replicated chromatin, an aberrant engagement of NHEJ factors, and HDAC1 and 2 as the only catalytic subunits of histone deacetylases ^29^. These features point to excessive deacetylase activity acting on replication-coupled chromatin in disease-relevant cells ^29^. We first asked whether depleting candidate deacetylases restores H4K16ac in these cells. HDAC2 knockdown increased the mean H4K16ac intensity by 37% relative to the non-targeting control, and SIRT1 knockdown increased the mean by 15% (Figure 5A).

**Figure 5.**
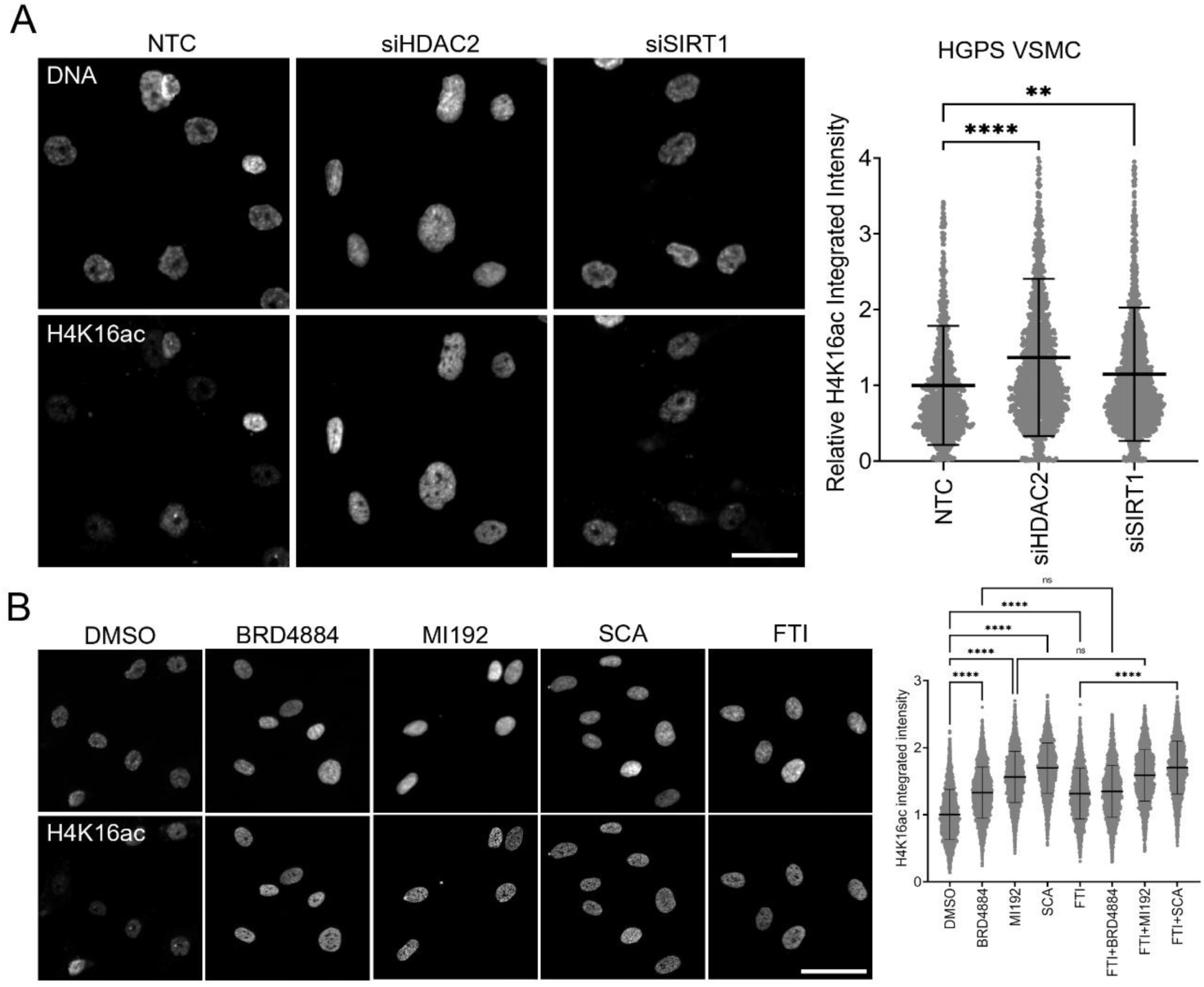
HDAC2 is the primary contributor to H4K16ac loss in HGPS VSMCs. (A) HGPS VSMCs transfected with siNTC, siHDAC2, or siSIRT1, with single-cell quantification of nuclear H4K16ac integrated intensity. Bars show the mean ± SD of three biological-replicate means. RM one-way ANOVA, Dunnett’s vs. NTC. Scale bar, 20 µm. (B) HGPS VSMCs treated for 24 h with DMSO, BRD4884, MI192, SCA, or lonafarnib (FTI), alone or in the indicated combinations. Representative images are shown for DMSO, BRD4884, MI192, SCA, and FTI. Bars show the mean ± SD of three biological-replicate means. RM one-way ANOVA, Dunnett’s vs. DMSO; prespecified FTI vs. FTI-combination comparisons, Šídák. ns, not significant; **p < 0.01; ****p < 0.0001. Scale bar, 20 µm.

Immunoblotting verified efficient target depletion in these cells (Figure S5A). This raises the possibility that HDAC2-specific inhibition may have therapeutic potential. We next evaluated HDAC inhibitors using working doses selected from titrations in HeLa cells (Figure S4A). After 24 h, SCA produced the largest increase in mean H4K16ac at 101%, MI192 increased the mean by 77%, and BRD4884 by 62% (Figure 5B). SCA gave the strongest response, followed by MI192 and BRD4884. Thus, in HGPS VSMCs, HDAC2 emerges as the principal H4K16 deacetylase and the most effective target for restoring H4K16ac, whereas SIRT1 targeting provides minimal benefit. Because farnesyltransferase inhibition remains the current therapeutic approach for HGPS, we next asked whether farnesyltransferase inhibitor (FTI) (Lonafarnib) altered H4K16ac either alone or in combination with HDAC2-directed inhibitors in HGPS VSMCs. FTI alone increased H4K16ac relative to DMSO by 39% (P<0.05). Combining FTI with BRD4884, MI192, or SCA maintained elevated H4K16ac levels, with combination-treated cells remaining among the highest H4K16ac values observed relative to DMSO-treated HGPS VSMCs (Figure 5B). Thus, farnesyltransferase inhibition was compatible with elevated H4K16ac in combination with HDAC2-directed inhibition.

### HDAC2-directed inhibition improves nuclear architecture, preserves proliferative capacity, and limits senescence-associated phenotypes in HGPS VSMCs

HGPS VSMCs display accelerated functional decline, consistent with atherogenesis observed in patients. This is characterized by atypical nuclear morphology, diminished proliferative capacity, and premature transition into cellular senescence ^29^, aligning with previous findings in this model ^42,43,73^. We therefore assessed whether inhibiting the H4K16ac eraser axis using these same HDAC2-biased compounds (BRD4884, MI192, SCA), alone and in combination with the FTI, could rescue these phenotypes while using the SIRT1 inhibitor EX527 and unaffected parental control VSMCs as comparators. High-content immunofluorescence microscopy was performed in asynchronous cultures. For nuclear architecture, we quantified form factor, eccentricity, and solidity. Relative to unaffected control VSMCs, HGPS DMSO nuclei were markedly dysmorphic across all metrics, and the HDAC2 inhibitors, alone or combined with FTI, restored form factor, eccentricity, and solidity toward control values. SCA typically showed the largest improvement in form factor and solidity, with BRD4884 closely comparable and MI192 slightly less effective, while eccentricity declined in parallel, indicating reduced elongation. FTI+SCA approached the unaffected control, whereas the SIRT1 inhibitor EX527 did not significantly improve nuclear morphology and FTI alone improved form factor only modestly (Figure 6A; form factor, eccentricity and solidity are plotted as separate panels, left to right). Quantitatively, HGPS-DMSO nuclei differed from parental controls in form factor (0.60 vs 0.83), eccentricity (0.70 vs 0.47), and solidity (0.84 vs 0.95); SCA restored these toward control (form factor 0.72, eccentricity 0.56, solidity 0.91; all P<0.01), BRD4884 and MI192 were comparable (form factor 0.68), and FTI+SCA gave the largest rescue (form factor 0.78, eccentricity 0.54, solidity 0.92; P≤0.001), whereas EX527 was not significant (form factor 0.61, eccentricity 0.72, solidity 0.84), and FTI alone produced a modest improvement in form factor (0.65, P<0.01) without significantly altering eccentricity (0.66) or solidity (0.86) (n=3 biological replicates; statistics computed on replicate means). Farnesyltransferase inhibition has repeatedly been reported to improve nuclear morphology in HGPS and ZMPSTE24-deficient patient fibroblasts ^35,74,75^. The limited FTI-alone effect here is therefore most plausibly attributable to cell type (VSMC rather than fibroblast), to the treatment window used, and to our use of unbiased continuous shape metrics across whole asynchronous populations rather than categorical scoring of blebbed nuclei; consistent with a real but smaller contribution, FTI augmented every HDAC2 inhibitor in combination.

**Figure 6.**
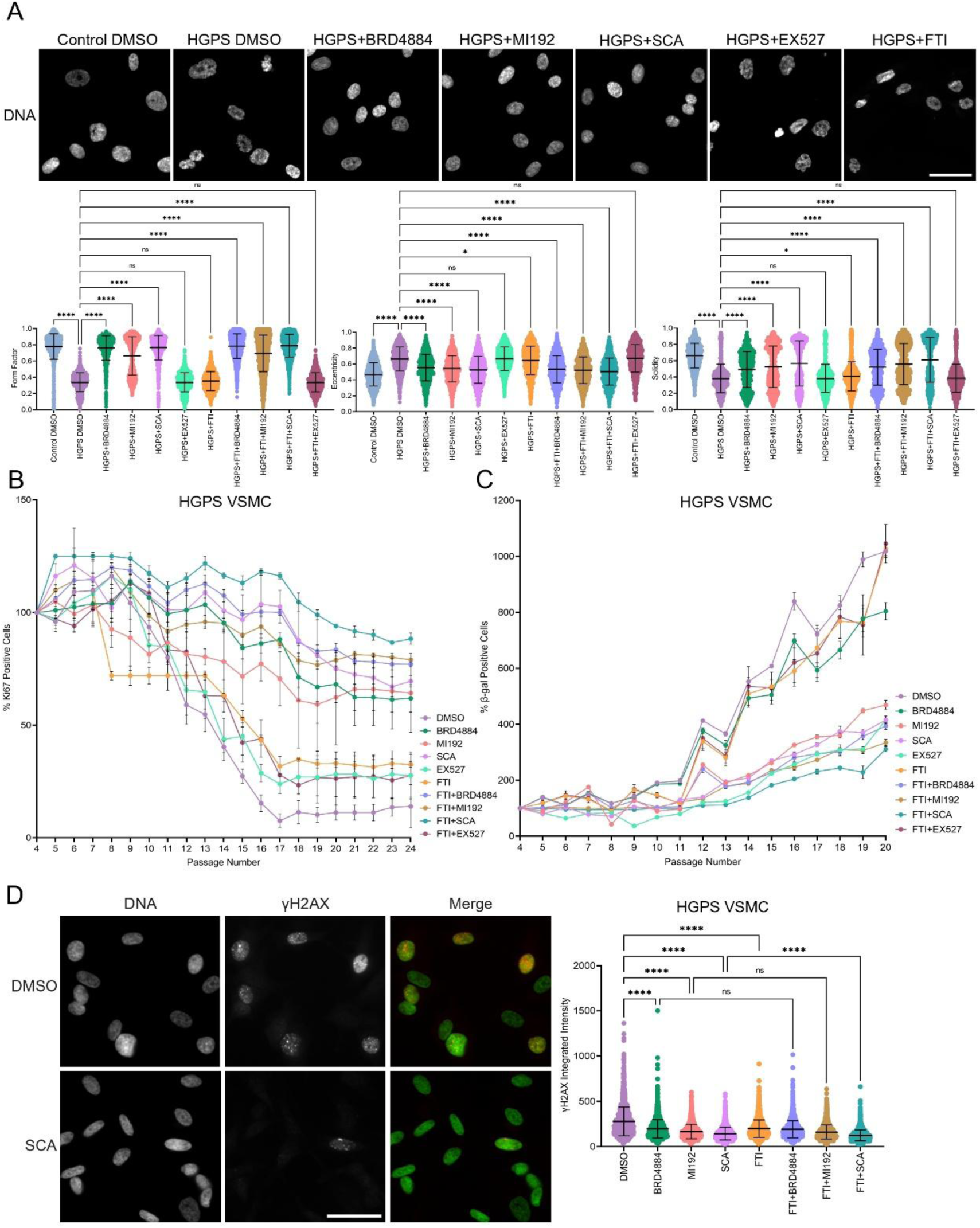
HDAC2 inhibition, alone and in combination with lonafarnib, improves nuclear morphology, preserves proliferation, and limits senescence and DNA-damage phenotypes in HGPS VSMCs. (A) Nuclear morphology in unaffected control and HGPS VSMCs treated with DMSO, BRD4884, MI192, SCA, EX527, or FTI, alone or in the indicated combinations, with representative Hoechst-stained images shown for selected conditions and quantification of form factor, eccentricity, and solidity. Bars show the mean ± SD of three biological-replicate means. Each parameter was analyzed separately: RM one-way ANOVA, Dunnett’s vs. DMSO-treated HGPS VSMCs; prespecified FTI vs. FTI-combination comparisons, Šídák. (B) Ki67 positivity during serial passage under the treatment conditions described in (A), normalised to the corresponding passage-4 value (set to 100%). Lines and error bars represent the mean ± SD from three independent biological replicates. Two-way RM ANOVA (treatment × passage) with Geisser–Greenhouse correction, Dunnett’s vs. DMSO-treated HGPS VSMCs. (C) SA-β-gal positivity during serial passage under the treatment conditions described in (A), normalised to the corresponding passage-4 value (set to 100%). Lines and error bars represent the mean ± SD from three independent biological replicates. Two-way RM ANOVA (treatment × passage) with Geisser– Greenhouse correction, Dunnett’s vs. DMSO-treated HGPS VSMCs. (D) Nuclear γH2AX integrated intensity in passage-14 HGPS VSMCs under the indicated single-agent and combination treatments, with representative DNA, γH2AX, and merged images shown for DMSO- and SCA-treated cells (merge: DNA, green; γH2AX, red). Bars show the mean ± SD of three biological-replicate means. RM one-way ANOVA, Dunnett’s vs. DMSO-treated HGPS VSMCs; prespecified FTI vs. FTI-combination comparisons, Šídák. ns, not significant; *p < 0.05; ****p < 0.0001. Scale bar for γH2AX images, 50 µm.

To evaluate proliferative capacity, we quantified Ki67 positivity in HGPS VSMCs across serial passages, normalised to the corresponding passage-4 value (set to 100%). In DMSO treated cultures, Ki67 positivity declined after passage 10 and continued to fall through late passages (Figure 6B). In contrast, class I HDAC inhibitors that show robust HDAC2 inhibition maintained higher Ki67 positivity across the same interval. BRD4884, MI192, and SCA each produced significantly higher Ki67 than DMSO from passage 11 onward, with P values less than 0.0001 from passage 17. The separation from vehicle widened as passages increased, with SCA typically retaining the highest Ki67 positivity while BRD4884 and MI192 also retained elevated Ki67 positivity. FTI alone partially preserved Ki67 relative to DMSO but remained below the HDAC2 inhibitors, whereas combining FTI with BRD4884, MI192, or SCA sustained the highest Ki67 positivity through passage 24, with FTI+SCA the most effective. EX527 tracked with DMSO and did not differ significantly at any passage in this range, whereas FTI+EX527 tracked FTI alone. By passage 20, Ki67 positivity was 12% in DMSO versus 89% in parental controls; HDAC2 inhibition preserved proliferation (SCA 75%, BRD4884 67%, MI192 63%), FTI combinations were highest (FTI+SCA 84%, FTI+BRD4884 76%, FTI+MI192 73%), FTI alone was intermediate (46%), and EX527 tracked DMSO (14%). These data indicate that HDAC2 inhibition functions as a senomorphic, preserving Ki67 positivity across late passages in HGPS VSMCs, whereas SIRT1 inhibition does not.

HGPS VSMCs are characterized by progressive accumulation of senescence-associated β-galactosidase (SA-β-gal) with passage ^29^. We therefore quantified SA-β-gal positivity across serial passages using the BioTracker 519 Green kit, normalised to the corresponding passage-4 value (set to 100%). Under DMSO treatment, β-gal–positive cells rose sharply after passage 12 (Figure 6C). HDAC2-directed inhibitors blunted this rise, with separation from DMSO evident at passage 12 and persisting through late passages. Averaged across passages, β-gal positivity was lower by 58% with BRD4884, 62% with MI192, and 73% with SCA (P<0.0001 for each). EX527 was 15% lower and not significant (P=0.39). These data demonstrate that HDAC2 inhibition reduces accumulation of the β-gal senescence marker in HGPS VSMCs, whereas SIRT1 inhibition does not under the conditions tested.

Phosphorylation of H2AX on Ser139 (γH2AX) marks chromatin flanking DNA double-strand breaks ^76^, and HGPS VSMCs accumulate DNA damage detectable as elevated γH2AX ^29^. Because restoring H4K16ac is predicted to relieve the NHEJ-biased, repair-deficient state of HGPS chromatin, we quantified γH2AX after 24 h of treatment. Relative to DMSO, the HDAC2 inhibitors reduced γH2AX integrated intensity, with SCA producing the largest reduction; FTI– inhibitor combinations further lowered γH2AX, with FTI+SCA reaching the lowest levels, whereas FTI alone was less effective (Figure 6D). Quantitatively, HGPS-DMSO γH2AX integrated intensity was 3.8-fold that of parental controls; relative to DMSO, SCA reduced γH2AX by 50%, MI192 by 32%, and BRD4884 by 26% (P<0.01), and FTI combinations further lowered it (FTI+SCA 60%, FTI+MI192 46%, FTI+BRD4884 45%; P<0.001), whereas EX527 (8%) was not significant and FTI alone produced a smaller but significant reduction (19%, P<0.001). Representative images confirmed a marked loss of γH2AX signal in SCA-treated relative to DMSO-treated cells (Figure 6D).

## Discussion

Building on our observation that H4K16ac is depleted in HGPS VSMCs ^29^, we sought to identify the enzymes that determine its steady-state abundance and to test whether inhibiting H4K16 deacetylation could delay senescence during serial passage. An siRNA screen targeting KAT5, KAT8, all eleven HDACs, and all seven sirtuins identified HDAC2, SIRT1, and KAT8 as the prominent regulators of H4K16ac in both HeLa and U2OS cells, with depletion of either HDAC2 or SIRT1 increasing per-nucleus H4K16ac by approximately 50%. HDAC2 was validated by imaging and immunoblotting of acid-extracted histones, whereas the SIRT1 effect was supported by imaging and reciprocal overexpression. The U2OS result is consistent with the original characterization of human SIRT1 as an H4K16 deacetylase, which paired in vitro deacetylation of H4K16 with RNAi-mediated SIRT1 depletion in U2OS cells ^52^. The HeLa result is harder to reconcile with the report that combined SIRT1 and SIRT2 depletion produces little or no increase in H4K16ac in HeLa cells ^32^. Our SIRT1 effect in HeLa rests on per-nucleus imaging and reciprocal overexpression rather than on immunoblotting of bulk histones, and an increase confined to a subset of nuclei or to a limited genomic compartment may be detected more readily by imaging than by immunoblot. Our findings are also consistent with previous reports that broad HDAC inhibition raises H4K16ac in Zmpste24-deficient progeroid cells ^26^ and that class I-selective inhibition with MS275 raises it in HGPS fibroblasts ^31^. However, the two screen hits were functionally distinct in HGPS VSMCs. HDAC2-directed inhibition attenuated γH2AX signaling, senescence-associated β-galactosidase accumulation, loss of Ki67 positivity, and abnormal nuclear morphology, whereas SIRT1 inhibition with EX527 neither raised H4K16ac effectively nor improved any phenotype tested, and SIRT1 depletion was markedly less effective at raising H4K16ac in HGPS VSMCs than in the cancer lines. SIRT1 is therefore a genuine H4K16 deacetylase in these highly proliferative cancer cell lines but is not the principal disease-relevant eraser in HGPS VSMCs.

Our conclusion differs from a previous report that lamin A/C binds and activates HDAC2, that progerin weakens this interaction, and that H4K16ac is consequently elevated in HGPS fibroblasts ^30,31^. Several observations support H4K16 hypoacetylation in progeroid and aged cells, including vascular smooth muscle. In Zmpste24-deficient cells, reduced H4K16ac is associated with impaired nuclear-matrix association of MOF/KAT8 without loss of total MOF protein ^26^. In HGPS VSMCs, acetylation turnover measurements resolve both reduced H4K16 acetylation and increased deacetylation, proximity ligation shows reduced KAT8 association with replicating DNA, and KAT8 depletion in control VSMCs lowers H4K16ac and increases 53BP1 ^29^. Class I HDAC inhibition with MS275 also raises H4K16ac in HGPS fibroblasts ^31^, demonstrating that residual class I deacetylase activity restrains the mark even in progerin-expressing fibroblasts. In the same study the pan-HDAC inhibitor trichostatin A did not raise H4K16ac in HGPS fibroblasts, an outcome the authors attributed to concurrent restoration of the lamin A/C–HDAC2 interaction ^31^. Because these compounds also modify the interaction they are used to interrogate, the MS275 result is the more direct indication that class I enzymes continue to deacetylate H4K16 in progerin-expressing cells; consistent with this, trichostatin A raises H4K16ac in HGPS VSMCs ^29^. Reduced H4K16ac is additionally observed in primary coronary artery smooth muscle cells from aged donors, which occurs in the presence of low levels of progerin ^29^.

Our results are consistent with a model linking H4K16ac restoration to the defective S-phase DNA-damage response in HGPS. We previously found enrichment of the NuRD HDAC1/2 complex and aberrant accumulation of 53BP1, Ku70/80, and DNA-PKcs on newly replicated chromatin in HGPS VSMCs ^29^. 53BP1 is recruited to damaged chromatin bivalently, through tandem-Tudor recognition of H4K20me2 and through a ubiquitination-dependent recruitment motif that reads RNF168-generated H2AK15ub ^77–79^. TIP60-mediated acetylation of H2AK15 antagonizes this ubiquitination-dependent arm ^62^. Acetylation of histone H4K16 weakens 53BP1 binding to H4K20me2 and shifts damaged chromatin toward BRCA1 and homologous recombination ^16^. In addition, pre-existing H4K16ac favours homologous recombination in S phase ^18^. KAT8 supplies the bulk of H4K16ac ^9,10,48^, and KAT8 additionally contributes to repair-pathway choice at damage sites ^17,80^. Beyond this role, KAT8 is required for efficient repair by both pathways ^81^. Excess HDAC2 activity could therefore reinforce an NHEJ-biased chromatin state and impair homologous recombination through 53BP1-mediated limitation of end resection, a mechanism established in BRCA1-deficient cells ^82^. Consistent with this direction, HDAC inhibition increases BRCA1 and decreases 53BP1 occupancy at damaged chromatin, the reciprocal of the shift produced by acetyltransferase deficiency ^16^. Consistent with a selective role in the NHEJ arm, HDAC1/2 depletion impairs non-homologous end joining and sustains DNA-damage signalling, yet does not markedly affect damage markers induced by camptothecin, which generates S-phase double-strand breaks that are repaired by homologous recombination ^32^. Although HDAC1 and HDAC2 share extensive sequence identity and frequently coexist in Sin3A, NuRD, and CoREST complexes, HDAC1- and HDAC2-containing complexes can perform context-dependent, non-redundant functions^32,83,84^. Previous work reported joint HDAC1/2 control of H4K16ac and showed that re-expression of HDAC1 could reverse H4K16 hyperacetylation ^32^; HDAC2 was not tested for reconstitution. Studies in oocytes and erythroblasts identified selective requirements for HDAC2, and in erythroblasts single HDAC2 knockdown was sufficient to increase H4K16ac ^56,85^. Our results extend this literature by assessing the two isoforms in intact somatic cells. Under our conditions, HDAC1 and HDAC2 were each depleted by more than 80%, yet only HDAC2 depletion increased H4K16ac in HeLa and U2OS cells. Conversely, EGFP-HDAC2 overexpression lowered H4K16ac, whereas EGFP-HDAC1 did not. This differs from earlier work, in which an increase was observed only after combined HDAC1/2 depletion ^32^. Notably their sufficiency experiment only tested re-expressed HDAC1 and was done in a depletion background, whereas ours compared overexpression of each isoform in otherwise intact cells. Collectively, the reciprocal loss- and gain-of-function data identify HDAC2 as the predominant class I H4K16 deacetylase in the systems tested. Isoform-biased perturbation is also achievable pharmacologically without relying on a single selective compound: BRD4884 inhibits HDAC1 and HDAC2 and MI192 inhibits HDAC2 and HDAC3 ^60,61^, so HDAC2 is the only isoform both engage at the concentrations used, and both raised H4K16ac.

Complementary evidence from the acetyltransferase arm shows that KAT8, the principal H4K16 acetyltransferase, is downregulated in aged human vessels and senescent endothelial cells and that its restoration attenuates vascular senescence, although H4K16ac was not measured in that study ^86^. Thus, increasing writer activity or limiting the dominant eraser converges on KAT8-dependent H4K16 acetylation as a candidate node in vascular aging. Short-term HDAC1/2 inhibition reduced renal fibrosis and altered age-associated transcriptional programs in the kidney, brain, and heart of aged mice ^87^, supporting in vivo evaluation of this strategy. We found that SCA combined with farnesyltransferase inhibition was well tolerated and produced greater effects on nuclear morphology and senescence markers than farnesyltransferase inhibition alone, which by itself improves nuclear morphology in progeroid patient cells ^74^; these experiments were not designed to distinguish additivity from synergy. This provides a rationale for preclinical evaluation of HDAC2-directed strategies in combination with farnesyltransferase inhibition in HGPS models.

Santacruzamate A (SCA) is a natural product with structural similarity to SAHA and was initially reported as a picomolar HDAC2 inhibitor with more than 3,500-fold selectivity over the class II enzymes HDAC4 and HDAC6; HDAC1, HDAC3 and HDAC8 were not tested^70^. However, the same group later could not reproduce this activity. In repeat assays, SCA was inactive against HDAC2 at 1 µM and inhibited all eleven HDAC isoforms only at 5–10 µM, and was inactive against sirtuins and DNA methyltransferases at those concentrations, arguing against potent or selective HDAC2 inhibition in vitro ^88^.

Despite the unresolved biochemical selectivity of SCA ^70,88^, several studies have reported cellular phenotypes concordant with genetic loss of HDAC2. In colorectal cancer cells, HDAC2 knockout increased H3K27ac-dependent recruitment of the BRD4–p-P65 complex to the NLRP3 promoter and enhanced NLRP3 transcription, and SCA reproduced the resulting pyroptotic and antitumour phenotype in combination with 5-fluorouracil or regorafenib, although the cellular concentration of SCA was not reported in that study ^89^. In a hepatocyte-injury model, Santacruzamate A (SCA; also known as CAY10683) at 120 nM produced effects on the mitochondrial apoptosis pathway concordant with HDAC2 knockdown and opposite to HDAC2 overexpression ^90^. Similar concordance was reported in porcine deltacoronavirus infection and in a colorectal-cancer metastasis model ^91,92^, although the concentrations used in those studies, 75 µM and 50 µM respectively, are ones at which broad HDAC inhibition and additional mechanisms are plausible ^88^. Separately, oral SCA reduced microglial activation in the anterior cingulate cortex of mice and lowered IL-1β, IL-6, and TNF-α in LPS-stimulated BV2 cells although this study implicated soluble epoxide hydrolase as a candidate target ^93^. However, that activity is weak and micromolar (IC50 7 µM with cellular effects at 10 µM) and does not account for the effects observed here at 10 nM ^93^. HDAC2 remains a preclinical therapeutic target in Alzheimer’s disease, although disease-modifying efficacy has not been established clinically ^94^.

Together, the reciprocal effects of HDAC2 depletion and overexpression in HeLa and U2OS cells, and of HDAC2 depletion and pharmacological inhibition in HGPS VSMCs, identify HDAC2 as the major class I determinant of steady-state H4K16ac in the models tested, with the inhibitor panel supporting its potential as a means to compensate for reduced KAT8 acetyltransferase activity. HDAC2 inhibition restores H4K16ac and attenuates nuclear shape alterations, γH2AX signaling, senescence-associated β-galactosidase accumulation, and loss of the Ki67-positive fraction. These findings establish HDAC2 as a candidate epigenetic target for mitigating vascular deterioration in HGPS and provide a rationale for isoform-biased inhibitor development and combination testing with farnesyltransferase inhibition.

## Limitations of the study

Our assignment of HDAC2 as the principal class I H4K16 deacetylase is bounded by the systems tested. The screen was performed in HeLa and U2OS cells and validated in iPSC-derived unaffected control and HGPS VSMCs; while reciprocal depletion and overexpression in the cancer lines, and depletion and pharmacological inhibition in the VSMC models, consistently support an HDAC2-preferential effect, they do not exclude HDAC1 activity at specific loci, cell-cycle stages, or corepressor complexes. “Principal” therefore denotes the dominant contributor to steady-state bulk H4K16ac in these models. We do not know whether it persists as the exclusive H4K16 class I deacetylase in all contexts. An earlier study reported H4K16 hyperacetylation only after combined HDAC1/2 depletion in U2OS cells ^32^, and we cannot exclude that differences in depletion efficiency or in readout account for the discrepancy with our single-isoform result in the same cell line.

Convergence between genetic perturbation, two benzamides with non-overlapping secondary targets, and the structurally distinct Santacruzamate A strengthens the assignment but does not exclude off-target contributions. Because HDAC2 acts on histone and non-histone substrates beyond H4K16, the observed improvements in nuclear morphology, γH2AX signaling, proliferative capacity, and SA-β-gal accumulation may not be due solely to H4K16ac restoration.

Finally, the disease-modifying experiments were confined to cultured iPSC-derived VSMCs and serial-passage assays, and so do not address in vivo efficacy, tissue specificity, pharmacokinetics, long-term safety, or improvement of vascular disease. Clinical experience with broad-spectrum HDAC inhibitors includes gastrointestinal, hematologic, and cardiac toxicities, some of which are dose-limiting ^65–68^. The lonafarnib experiments establish compatibility and absence of detectable antagonism but were not designed to determine synergy or define optimal dosing. Validation in additional cellular backgrounds and *in vivo* HGPS models will be required to assess the therapeutic potential and safety of sustained HDAC2 inhibition.

## Resource availability

### Lead contact

Further information and requests for resources and reagents should be directed to and will be fulfilled by the lead contact, Michael J. Hendzel.

### Materials availability

All siRNAs and small-molecule inhibitors used in this study were obtained commercially; they are not available for distribution from the lead contact. Plasmids and cell lines are available as set out below.

The induced pluripotent stem cell lines from which the vascular smooth muscle cells were differentiated were described previously ^73^ and were characterized in the vascular model used here ^29^. The HGPS iPSC lines used in this study were 0031C, 1671Q, and AG1B; control iPSC lines were 0901C, 168D2, and BJ1C, with lines 0901C and 168D2, representing lines derived from fibroblasts from the parents of 1671Q. Lines 0031C, 0901C, 1671Q and 168D2 are available from the Progeria Research Foundation under a Materials Transfer Agreement. BJ1C and AG1B are available directly from W.L. Stanford. No new cell lines were generated in the present study.

The GFP-tagged HDAC1, HDAC2 and HDAC3 expression constructs used for the overexpression experiments are available from the lead contact on request. The SIRT1 expression plasmid is available from Addgene (plasmid #13812).

### Data and code availability

All data reported in this paper will be shared by the lead contact upon request.

This paper does not report original code.

Any additional information required to reanalyze the data reported in this paper is available from the lead contact upon request.

## Experimental model and study participant details

### Immortalized human cell lines

HeLa cells (female origin) were used for the siRNA screen of KAT5, KAT8, HDAC1 through HDAC11 and SIRT1 through SIRT7, for the HDAC and SIRT1 overexpression experiments, and for the inhibitor dose titrations from which working concentrations were selected. U2OS cells (female origin) were used as an independent human cell line to test whether the H4K16ac dependencies identified in HeLa generalize beyond a single background. Both lines were grown in High Glucose DMEM (Gibco) supplemented with 10% fetal bovine serum (Gibco) and 1% penicillin-streptomycin (Thermo Fisher) at 37°C in a humidified atmosphere containing 5% CO₂ and were screened routinely for mycoplasma contamination.

### Human iPSC-derived vascular smooth muscle cells

Vascular smooth muscle cells were differentiated from human induced pluripotent stem cell lines reprogrammed from fibroblasts obtained from patients with Hutchinson-Gilford progeria syndrome and from unaffected control individuals. These lines were described previously ^73^ and the vascular model was characterized previously ^29^. Three HGPS lines (0031C, 1671Q, and AG1B) and a combination of parental (0901C, 168D2) and unrelated (BJ1C) control iPSC lines were used. Because reprogramming resets the progeroid epigenome and undifferentiated iPSCs do not express lamin A, progerin-associated phenotypes emerge only after differentiation and accumulate with passage ^73^, making this model suitable for following the onset and progression of senescence.

Undifferentiated iPSCs were maintained on Matrigel-coated plates in E8 medium. Differentiation followed an embryoid body protocol as modified previously ^29^: colonies were dissociated with collagenase IV, embryoid bodies were formed in low-attachment plates for 7 to 10 days, plated on 0.1% gelatin-coated plates for 3 to 5 days, and outgrowths were replated on Matrigel in Medium 231 with growth supplement. For differentiation to vascular smooth muscle cells, cells were plated on gelatin-coated plates in Medium 231 with smooth muscle differentiation supplement (Thermo Fisher) for 7 days and characterized at each passage.

Because progerin accumulation, nuclear dysmorphia, DNA-damage signaling and senescence in this model are passage-dependent, single-timepoint assays were performed on passage 14 cultures, and the senescence time course (Ki67 and senescence-associated β-galactosidase) was followed from passage 4 through passage 24. Parental control VSMCs were carried in parallel at every passage as the unaffected comparator.

## Method details

### Pharmacological Treatments

To examine the role of HDAC inhibition in modulating histone acetylation and associated cellular phenotypes, cells were treated with BRD4884 (2 µM), MI192 (1 µM), EX527 (10 µM), trichostatin A (TSA; 1.5 µM), Quisinostat (JNJ-26481585; QS; 250 nM), Santacruzamate A (SCA; 10 nM), or lonafarnib (FTI; 3 µM). Cells were incubated with each compound for 24 hours prior to fixation, protein extraction, or imaging. For combination treatments, lonafarnib was co-administered with BRD4884, MI192, SCA, or EX527 at the concentrations indicated above.

To determine the optimal concentration for each compound, a titration series was performed in which increasing doses were tested for their effects on proliferation, cell cycle distribution, and histone acetylation. Proliferation was monitored by seeding equal numbers of cells and measuring population doublings over time. In parallel, H4K16ac levels were assessed by both immunofluorescence and western blot analysis to identify the dose that maximally increased H4K16 acetylation without reducing proliferation or altering cell cycle distribution. Cell cycle profiles were evaluated by quantifying DNA content using DAPI fluorescence intensity. Final treatment concentrations were chosen based on their capacity to enhance H4K16ac while maintaining proliferative capacity and normal cell cycle profiles.

### siRNA Knockdown and Transfection

For systematic depletion of HDACs and sirtuins, HeLa and U2OS cells were transfected with ON-TARGETplus SMARTpool siRNAs (Dharmacon) targeting HDAC1–HDAC11 and SIRT1– SIRT7. Selected candidates identified from this screen were subsequently depleted in unaffected control and HGPS VSMCs. A non-targeting siRNA (NT-siRNA) was used as a negative control, and Lamin A/C siRNA served as a positive control. Transfections were carried out using RNAiMAX (Invitrogen) in Opti-MEM (Gibco) at a final siRNA concentration of 40 nM. Cells were processed 72 hours post-transfection for immunofluorescence, western blotting, or quantitative imaging analyses.

For HDAC overexpression experiments, HeLa cells were transfected with GFP-tagged constructs encoding HDAC1, HDAC2, HDAC3, or SIRT1 using Effectene (Qiagen) following the manufacturer’s guidelines. The SIRT1 plasmid was obtained from Addgene (#13812). Transfections were performed in cells seeded to 60–70% confluency in 6-well plates containing glass coverslips, using 0.4 µg of plasmid DNA per well. Eighteen hours post-transfection, cells were either fixed for immunofluorescence or lysed for protein extraction. Transfection efficiency was confirmed by GFP fluorescence and immunoblot analysis.

### Immunofluorescence and Imaging

Cells grown on glass coverslips were treated as described, fixed in 4% paraformaldehyde for 10 minutes at room temperature, permeabilized with 0.5% Triton X-100 in PBS for 10 minutes, and blocked in 3% bovine serum albumin (BSA) in PBS for 30 minutes. Cells were incubated with the following primary antibodies for 1 hour at room temperature: H4K16ac (Sigma-Aldrich, 07-329), Ki67 (Abcam, ab15580), Lamin A/C (Cell Signaling, 4777S), HDAC2 (Cell Signaling, 25458), SIRT1 (Abcam, ab32441), and γH2AX (phospho-histone H2A.X Ser139). Following washes, cells were incubated with Alexa Fluor-conjugated secondary antibodies (Thermo Fisher) for 1 hour at room temperature and counterstained with Hoechst 33342 (1 µg/mL, Invitrogen). Coverslips were mounted in Mowiol and imaged.

High-content imaging for siRNA screens and nuclear morphology analyses was performed using a MetaXpress XLS system high-content analysis system (Molecular Devices, San Jose, CA, USA) equipped with a 20X/0.75 NA Plan-Apochromat objective and an sCMOS camera (2180×2180 pixels, 6.5 µm pixel size). For HDAC overexpression and pharmacological experiments, imaging was conducted using a Zeiss AxioImager microscope with a 40X/1.4 NA oil immersion objective and a Photometrics PRIME BSI camera.

### Image Analysis

Quantitative image analysis was performed using CellProfiler 4.2.8 (Broad Institute) ^95^ and ImageJ (NIH) ^96^. Nuclear morphology was assessed using automated segmentation to extract shape descriptors including form factor, eccentricity, and solidity. Ki67 positivity was calculated as the percentage of nuclei classified as Ki67-positive ^97^. Nuclear H4K16ac levels were measured as per-nucleus integrated fluorescence intensity following background correction. Ki67 and SA-β-gal positivity were determined from per-nucleus fluorescence intensities using a fixed intensity threshold that was applied consistently across all conditions and passages within each assay. These descriptors were defined as form factor (4π × area/perimeter²; circularity), eccentricity (eccentricity of the fitted ellipse; elongation) and solidity (area/convex-hull area; contour integrity). For the overexpression experiments, nuclear H4K16ac was quantified only in EGFP-positive nuclei, identified using a fixed EGFP intensity threshold applied consistently across all conditions. Nuclear γH2AX was quantified as per-nucleus integrated fluorescence intensity after background correction, using the same nuclear segmentation as for H4K16ac.

### Histone Extraction and Western Blotting

To quantify protein levels and histone acetylation, whole-cell proteins and histones were prepared separately. For analysis of histone acetylation, histones were isolated by acid extraction. Nuclear pellets were resuspended in ice-cold 0.2 M H₂SO₄ (0.4 N) and incubated with rotation for 2 hours at 4°C. Samples were centrifuged at 3,400 × g for 5 minutes at 4°C, and histones in the supernatant were precipitated with trichloroacetic acid (TCA) to a final concentration of 33% for 1 hour on ice. Precipitated histones were collected by centrifugation at 3,400 × g for 5 minutes at 4°C, washed once with ice-cold acetone containing 0.1% HCl and once with ice-cold 100% acetone, air-dried, and resuspended in ddH₂O. Equal amounts of acid-extracted histones (5 µg/lane) were resolved by 15% SDS-PAGE and transferred to nitrocellulose membranes (Bio-Rad) at 30 mA overnight at 4°C. Membranes were blocked in 5% non-fat milk in PBS for 1 hour and incubated overnight at 4°C with primary antibodies against H4K16ac and total histone H3. Following three TBST washes, membranes were incubated with IRDye-conjugated secondary antibodies (LI-COR, 1:5000) for 2 hours at room temperature. Fluorescence signals were imaged using the Odyssey Fc Imaging System (LI-COR), and band intensities were quantified using ImageJ. H4K16ac signal was normalized to total histone H3 (Sigma-Aldrich, 05-499), and values were expressed relative to the corresponding control condition (set to 1.0).

For analysis of non-histone proteins, cells were lysed in 2× SDS buffer (2% SDS, 50 mM Tris, pH 7.5, 10% glycerol, supplemented with protease inhibitors), sonicated for 1 minute (Model 705 Sonic Dismembrator, Fisher Scientific), and heated to 95°C for 10 minutes. Equal protein amounts (20 µg/lane) were resolved by SDS-PAGE and transferred to nitrocellulose membranes (Bio-Rad) at 30 mA overnight at 4°C. Membranes were blocked in 5% non-fat milk in PBS for 1 hour and incubated overnight at 4°C with primary antibodies against the indicated proteins, including HDAC2 and SIRT1, with GAPDH used as the loading control. Following three TBST washes, membranes were incubated with IRDye-conjugated secondary antibodies (LI-COR, 1:5000) for 2 hours at room temperature. Fluorescence signals were imaged using the Odyssey Fc Imaging System (LI-COR), and band intensities were quantified using ImageJ. Non-histone protein levels were normalized to GAPDH and expressed relative to the corresponding control condition.

### Proliferation and Senescence-Associated β-Galactosidase Assays

To measure proliferative capacity, equal numbers of cells were seeded, treated with inhibitors for 24 hours, and allowed to expand until reaching approximately 80% confluence. Cells were then trypsinized, counted using the Countess II Automated Cell Counter (Invitrogen), and reseeded at equal density. Population doublings (PD) at each passage were calculated as PD = log_2_ (N_h_/N_s_), where N_h_ represents the number of cells harvested and N_s_ represents the number of cells seeded. Cumulative population doublings were calculated by summing the PD values across successive passages. Ki67 positivity was assessed by immunofluorescence as described above and quantified as the percentage of nuclei classified as Ki67-positive at each passage. Serial-passage values were normalised to the corresponding passage-4 value, set to 100%.

Senescence-associated β-galactosidase (SA-βgal) activity was measured using the BioTracker 519 Green β-Gal Detection Kit (Sigma-Aldrich). The fluorescent β-gal substrate was diluted in culture medium to a final concentration of 1 µM, and cells were incubated for 15 minutes at 37°C in 5% CO_2._ Following staining, cells were washed two to three times with HBSS and imaged using the MetaXpress XLS high-content analysis system (Molecular Devices). The percentage of SA-βgal-positive cells was quantified from multiple fields of view for each biological replicate. Serial-passage values were normalised to the corresponding passage-4 value, set to 100%.

### Quantification and statistical analysis

Statistical analyses were performed using GraphPad Prism. For image-based assays, measurements from individual nuclei were averaged within each independent biological replicate before statistical analysis; individual nuclei were not treated as independent biological observations. The number of nuclei analyzed per condition is specified in the corresponding figure legends. Unless otherwise indicated, n = 3 independent biological replicates per condition. Data are presented as mean ± SD. Within each independent experiment, per-nucleus values were normalized to the mean of the corresponding control condition (NTC, EGFP-only or DMSO), which was set to 1; immunoblot band intensities were normalized to the appropriate loading control (total histone H3 for acid-extracted histones, GAPDH for whole-cell lysates) and expressed relative to the matched control condition. Dose-response data are expressed as a percentage of the corresponding DMSO-treated control. Where single-cell data are shown, violin plots or grey points display pooled single-cell measurements to illustrate the distribution and cell-to-cell variability within each condition; all statistical comparisons were performed on biological-replicate means.

For experiments in which treatment conditions were assayed in parallel within the same independent experiment, biological replicate was treated as a matched factor. Single-factor experiments containing three or more matched conditions were analyzed by repeated-measures one-way ANOVA followed by Dunnett’s multiple-comparisons test against the prespecified control condition. Pairwise immunoblot validation experiments were analyzed using two-tailed paired t-tests.

Serial-passage Ki67 and SA-β-gal data were analyzed by two-way repeated-measures ANOVA with treatment and passage as factors, followed by Dunnett’s multiple-comparisons test comparing each treatment with DMSO-treated HGPS VSMCs at the indicated passages. The Geisser–Greenhouse correction was applied to account for departures from sphericity.

For experiments evaluating lonafarnib-containing combinations, comparisons of individual treatments with DMSO-treated HGPS VSMCs were performed using Dunnett’s multiple-comparisons test, whereas prespecified comparisons between lonafarnib alone and the corresponding lonafarnib-containing combination treatments were evaluated using Šídák-adjusted multiple-comparisons tests.

The statistical test used for each experiment is specified in the corresponding figure legend. Statistical significance was defined as p < 0.05, and only statistically significant differences are indicated in the figures. Significance is indicated as ns, not significant; *p < 0.05; **p < 0.01; ***p < 0.001; and ****p < 0.0001.

## Acknowledgments

This work was supported by a CIHR project grant [PJT-178364] awarded to W.L.S. and M.J.H. We thank the Cell Imaging Facility at the Cross Cancer Institute for providing access to instrumentation and technical advice and support.

## Author contributions

M.J.H. and W.L.S. conceived the project with support from R.K. and M.N. M.J.H. and W.L.S. designed the methodology. Original drafting was done by R.K. R.K., M.N., M.J.H. and W.L.S., all contributed writing to the final document and were all involved in editing.

## Declaration of interests

The authors declare no competing interests.

## Declaration of generative AI and AI-assisted technologies in the manuscript preparation process

Claude and ChatGPT were used for rewording the text of the document for clarity and conciseness. ChatGPT Deep Research together with adversarial review by Claude was used to improve the accuracy of language when describing the published literature as well as identify any missing key references, including contradictory research breadth. The authors have reviewed and approved any edited text and take full responsibility for the content.

## Notes

### Competing Interest Statement

The authors have declared no competing interest.

### Summary of Updates

This revision expands the experimental analysis and refines the interpretation of HDAC2-dependent H4K16ac regulation in HGPS vascular smooth muscle cells. The title has been revised to HDAC2 inhibition restores H4K16ac and delays senescence in HGPS VSMCs in order to reflect the experimental system and the observed delay in senescence-associated phenotypes. New experiments evaluate lonafarnib alone and in combination with HDAC2-directed inhibitors, extending the analysis of H4K16ac restoration, nuclear morphology, Ki67 positivity and senescence-associated β-galactosidase accumulation. Figure 6 now includes nuclear γH2AX measurements and representative images assessing DNA-damage-associated signalling; these replace the H4K20me3 panel in the original version. An additional immunoblot analysis in Figure S2 distinguishes the reduction in H4K16ac fluorescence after HDAC3 depletion from the absence of a significant reduction in bulk histone H4K16ac. The statistical analysis section and figure legends have been revised to specify independent biological replicates, matched and repeated-measures analyses, multiple-comparison procedures and the meaning of single-cell distributions. Serial-passage Ki67 and β-galactosidase values are explicitly described as normalised to the corresponding passage-4 baseline, set to 100%. Methods have been revised and expanded, including cell-line provenance, treatment conditions, imaging and assay procedures, with additional resource, data-availability and author declarations. The Introduction and Discussion have been substantially revised to address context-dependent H4K16ac changes in aging, differences between HGPS fibroblasts and VSMCs, and contrasting reports concerning HDAC1/2 and SIRT1. The pharmacological interpretation now addresses uncertainty in Santacruzamate A selectivity and distinguishes combination effects from demonstrated synergy. A dedicated limitations section defines the scope of the conclusions and the need for further validation. References, figure legends, cross-references and terminology have also been updated. The central conclusion remains that HDAC2 is a predominant class I regulator of steady-state H4K16ac in the models tested and a candidate target for limiting senescence-associated phenotypes in HGPS VSMCs.

