## Supplemental Data for "HDAC2 inhibition restores H4K16ac and delays senescence in HGPS VSMCs"

### Figure S1

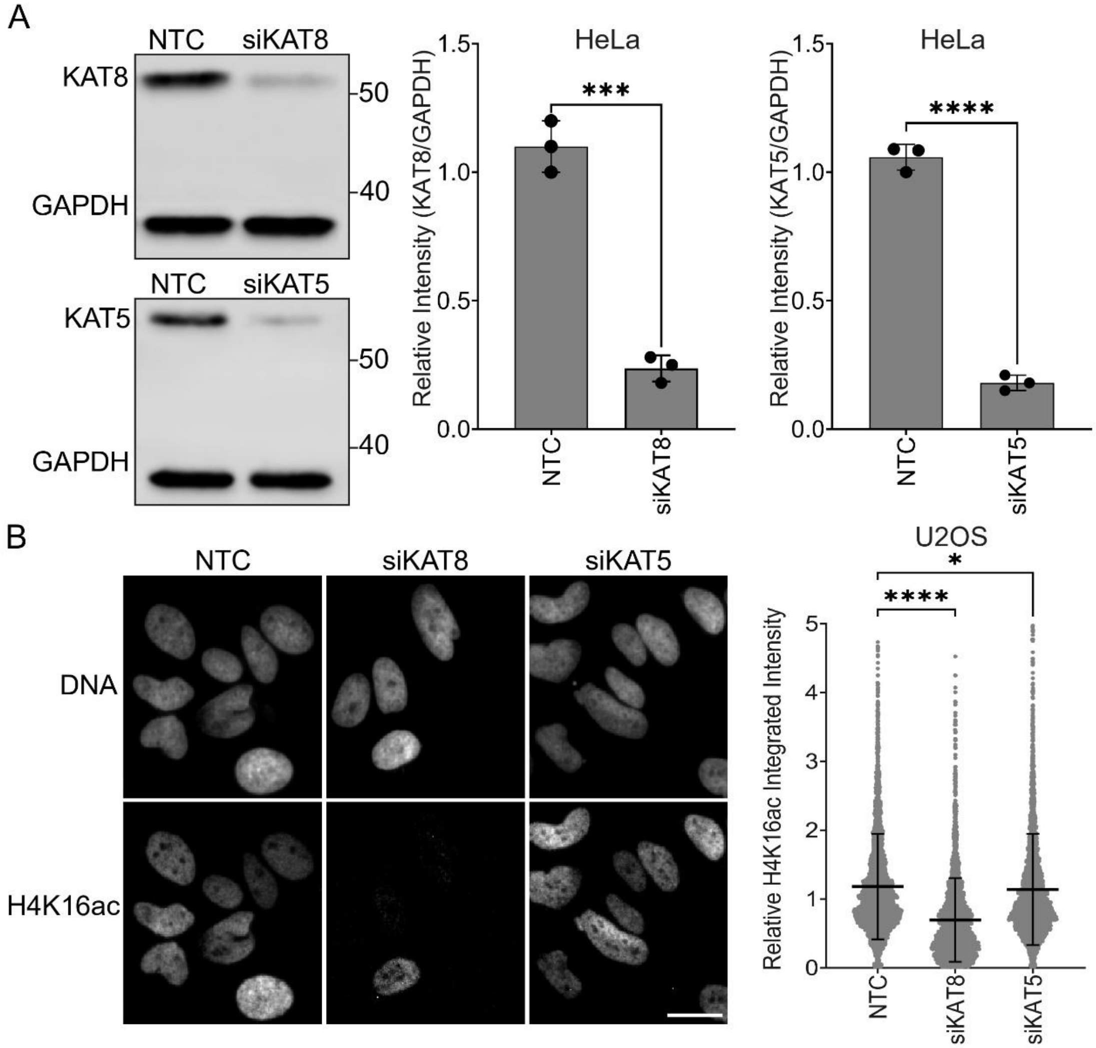


**Figure S1.** Validation of KAT8 and KAT5 depletion and KAT8-dependent H4K16 acetylation in U2OS cells, related to Figure 1. (A) Validation of KAT8 and KAT5 knockdown in HeLa cells transfected with siNTC, siKAT8, or siKAT5. Each point represents one independent biological replicate, and bars represent the mean ± SD from three independent biological replicates. Two-tailed paired t-test, each knockdown vs. matched NTC. (B) U2OS cells transfected with siNTC, siKAT8, or siKAT5, with representative images and single-cell quantification of nuclear H4K16ac integrated intensity. Approximately 1,000 cells were analyzed per condition per biological replicate across three independent biological replicates. RM one-way ANOVA, Dunnett's vs. NTC. Scale bar, 20 µm.

### Figure S2

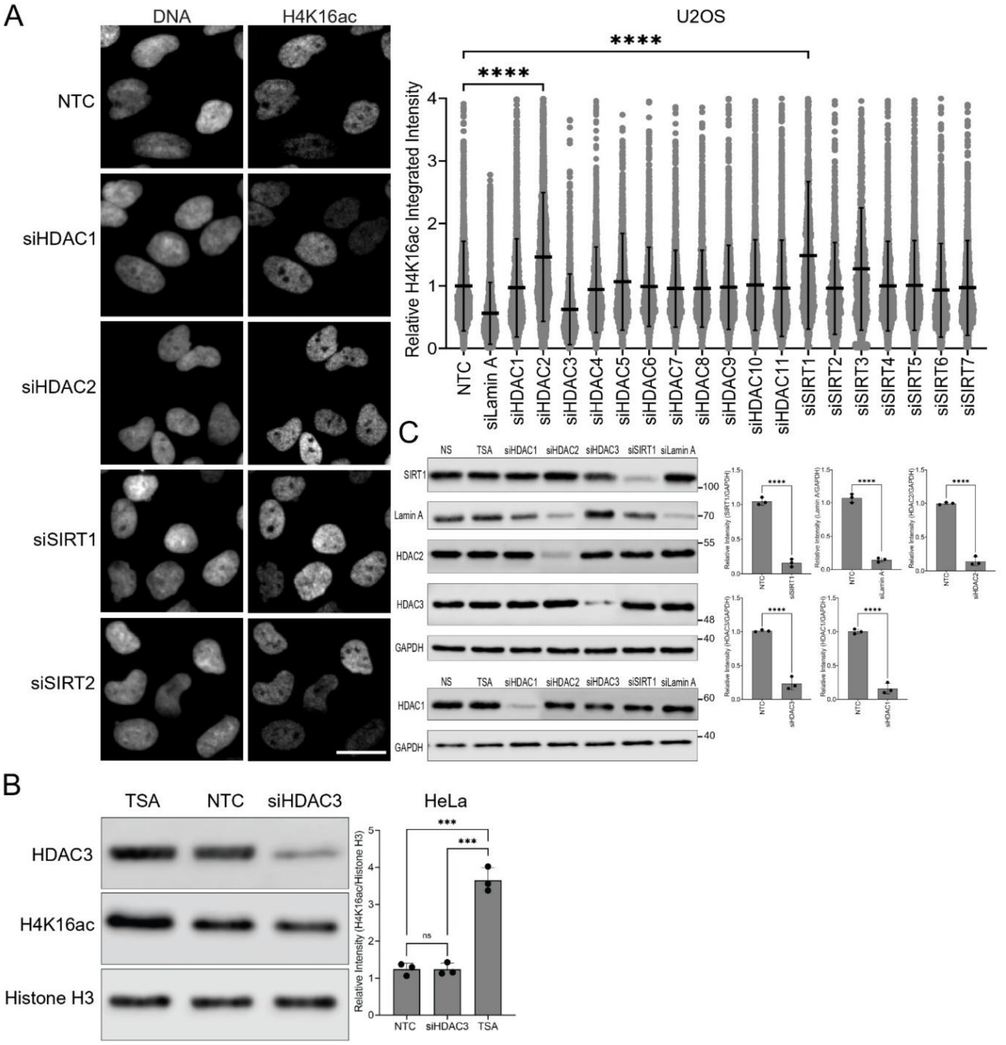


**Figure S2.** Validation of HDAC2-, SIRT1-, and HDAC3-dependent effects on steady-state H4K16ac, related to Figure 2. (A) U2OS cells transfected with siNTC or siRNAs targeting Lamin A, HDAC1–11, or SIRT1–7, with representative images shown for NTC, siHDAC1, siHDAC2, siSIRT1, and siSIRT2 and single-cell quantification of nuclear H4K16ac integrated intensity. Bars show the mean ± SD of three biological-replicate means, with at least 500 cells analyzed per condition per replicate. RM one-way ANOVA, Dunnett's vs. NTC. Scale bar, 20 µm. (B) Bulk H4K16ac in HeLa cells transfected with NTC or siHDAC3, with TSA included as a positive control for increased histone acetylation. Each point represents one independent biological replicate, and bars represent the mean ± SD from three independent biological replicates. RM one-way ANOVA, Dunnett's vs. NTC. (C) Validation of siRNA-mediated depletion in HeLa cells transfected with NTC, siHDAC1, siHDAC2, siHDAC3, siSIRT1, or siLamin A, with TSA-treated cells included as an additional control. Each point represents one independent biological replicate, and bars represent the mean ± SD from three independent biological replicates. Two-tailed paired t-test, each knockdown vs. matched NTC.

**Figure S3**

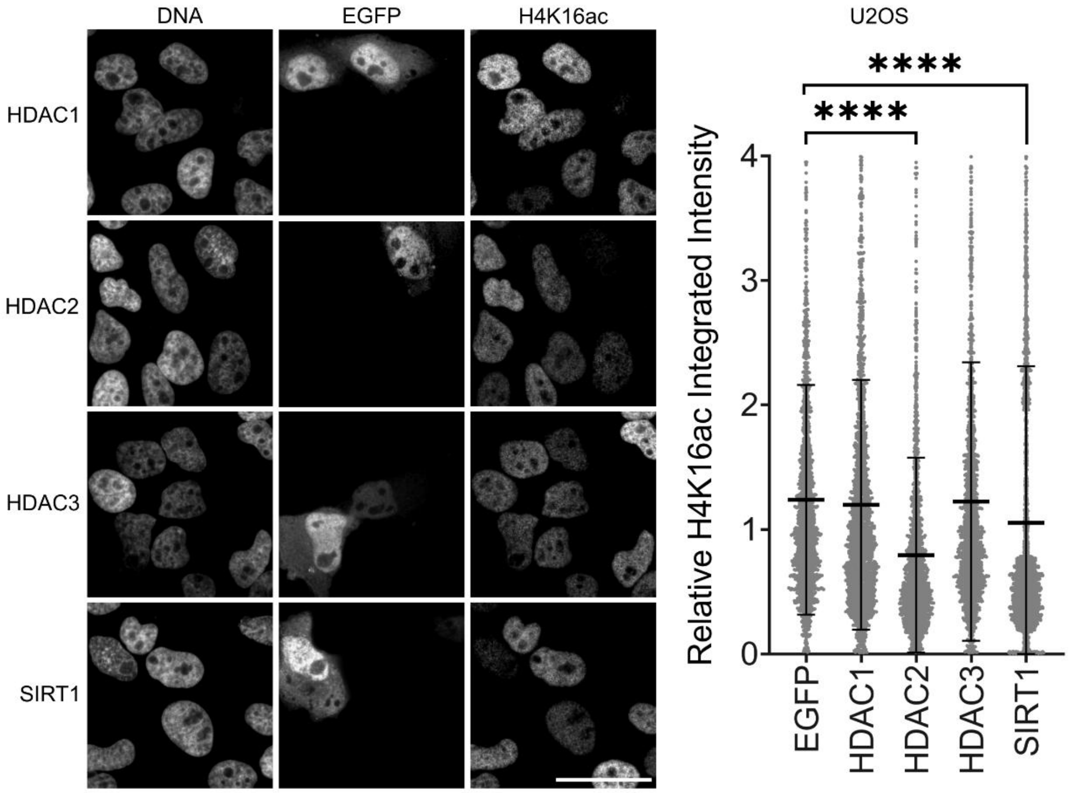


**Figure S3.** Overexpression of HDAC2 and SIRT1 reduces steady-state H4K16ac in U2OS cells, related to Figure 3. U2OS cells transfected with EGFP alone or EGFP-tagged HDAC1, HDAC2, HDAC3, or SIRT1, with representative images and quantification of nuclear H4K16ac integrated intensity in EGFP-positive cells. Bars show the mean ± SD of three biological-replicate means. RM one-way ANOVA, Dunnett's vs. the EGFP-only control. Scale bar, 10 μm.

### Figure S4

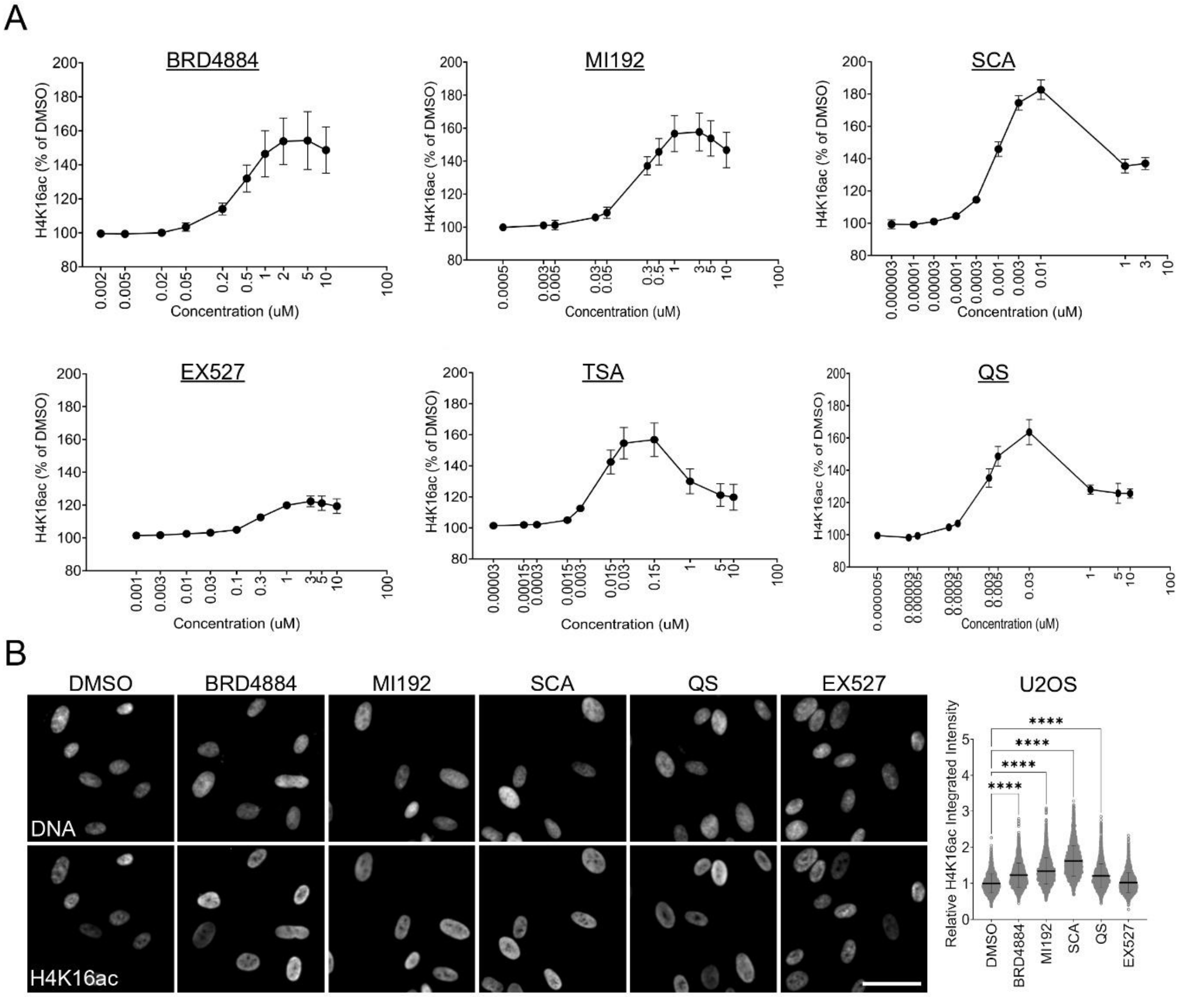


**Figure S4.** Dose-response optimization of H4K16ac-modulating compounds in HeLa cells and validation of responses in U2OS cells, related to Figure 4. (A) Dose-response analysis of BRD4884, MI192, SCA, EX527, TSA, and QS in HeLa cells treated for 24 h with the indicated concentrations. Points and error bars represent the mean ± SD from three independent biological replicates. (B) U2OS cells treated for 24 h with DMSO, BRD4884, MI192, SCA, QS, or EX527, with representative images and single-cell quantification of nuclear H4K16ac integrated intensity. Bars show the mean ± SD of three biological-replicate means. RM one-way ANOVA, Dunnett's vs. DMSO. Scale bar, 20 µm.

**Figure S5**

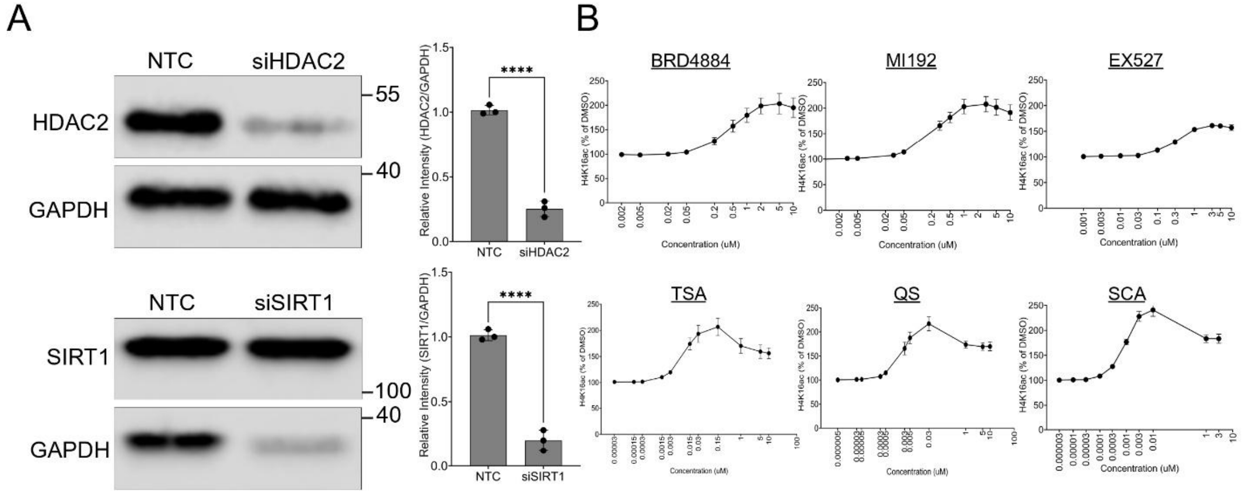


**Figure S5.** Validation of HDAC2 and SIRT1 depletion and compound dose-response analysis in HGPS VSMCs, related to Figure 5. (A) Validation of siRNA-mediated HDAC2 and SIRT1 depletion in HGPS VSMCs transfected with siNTC, siHDAC2, or siSIRT1. Each point represents one independent biological replicate, and bars represent the mean ± SD from three independent biological replicates. Two-tailed paired t-test, each knockdown vs. matched NTC. (B) Dose-response analysis of BRD4884, MI192, EX527, TSA, QS, and Santacruzamate A (SCA) in HGPS VSMCs treated for 24 h with the indicated concentrations. Points and error bars represent the mean ± SD from three independent biological replicates.
